# The Shape of Control: How Going in Circles Keeps RNA Catalytically Switched On

**DOI:** 10.64898/2026.01.28.702370

**Authors:** M. J. Billings, A. K. Chawla, A. Mulla-Feroze, A. M. Kietrys

## Abstract

RNase P was one of the first enzymes discovered to have an RNA-based catalytic component. Since its identification, it has been extensively studied, particularly in *E. coli*, due to the ability of its M1 RNA to exhibit *in vitro* catalytic activity even in the absence of associated protein. In this study, we report G-quadruplex formation as a potential regulatory mechanism that modulates the catalytic activity of this RNA. We observed a significantly higher propensity for G-quadruplex formation in the linear isoform (linM1) compared to its circular counterpart (circM1). G-quadruplex formation was confirmed through circular dichroism spectroscopy and a fluorescence-based assay using a G-quadruplex-binding small molecule. We compared the catalytic activity of linM1 and circM1 in lithium and potassium environments and found that G-quadruplex formation specifically reduced linM1 activity. Furthermore, we observed distinct condensate properties of linM1 in the presence or absence of G-quadruplex structures. Overall, our findings suggest that G-quadruplex formation serves as a regulatory switch for RNA activity in linM1, whereas circM1 resists G-quadruplex formation and remains catalytically active even under conditions that favor G-quadruplex assembly.

## Introduction

M1 RNA is a well-studied ribozyme, the core component of the pre-tRNA maturing bacterial RNase P enzyme^1^. While human RNase P has been shown to fold into G-quadruplexes^2^, no such counterpart study has been performed to evaluate G-quadruplex forming propensity of M1. In our previous study^3^, we reported a circular variant of M1 RNA (circM1). While circular DNA have been shown to have a lower G-quadruplex forming propensity^4^, circRNA, however, have not been investigated similarly for their comparative G-quadruplex forming propensity with respect to linear counterparts. Furthermore, while G-quadruplexes have been identified in the bacterial genome and the transcriptome^5^, they have not been shown to have any functional or regulatory role. Interestingly, RNA G-quadruplexes have been shown to facilitate condensation^6^, a process known to drive enzymatic reactions such as that of M1 RNA^7^.

### Evaluation of G-quadruplex in circular RNase P RNA isoforms

As we set forth on investigating the circM1 activity, we noticed a G-rich sequence. We utilized three tools, QGRS Mapper^8^, G4hunter^9^ and pqsfinder^10^ to probe for G-quadruplexes and found 5 putative G4 sequences (Fig. S1). To start off our investigation, we focused on the G4 with the highest G-score consisting of 5 tetrads and 3 bulging making it an extremely stable G4 (Figure S2). Firstly, we confirmed the presence of a parallel G-quadruplex in this sequence using circular dichroism as evident in the increased intensity of 260 nm peak in K^+^ environment, an established G4 stabilizing ion^11^ (Fig. 1D). Since it is not common for G4s to fold in Li^+^, we verified the absence of any K^+^ ions in the Li^+^ folding buffer using ESI-MS (data not shown). However, CD could not be utilized to compare the circular and linear isoforms of M1 due to the presence of other interfering structures in the full length transcript^12,13^ (Fig. S3).

**Figure 1.**
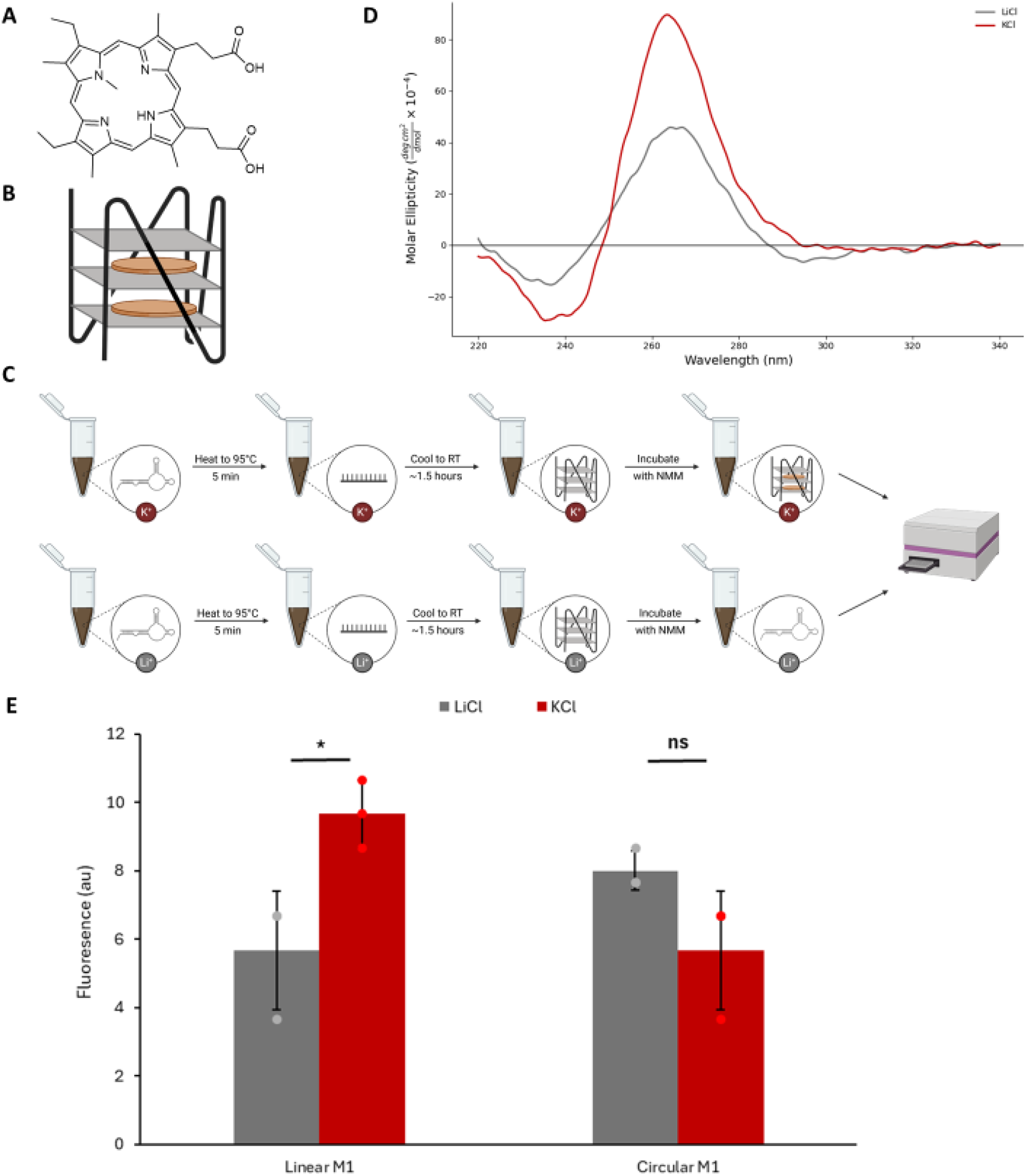
Linear M1 RNA demonstrates higher G4-forming propensity compared to circular isoform. (A) Chemical structure of N-methyl mesoporphyrin IX (NMM) (B) NMM stacks on pre-formed G-quadruplexes enhancing their stability (C) Schematic of NMM fluorescence assay (D) Circular dichroism spectra supporting G4 formation in predicted 47nt G4 M1 fragment (E) Higher NMM fluorescence for linM1 in the presence of K^+^ ions indicating G4-formation. No significant difference in fluorescence observed for circM1. For quantitative analysis, n=3. Student’s t-test were performed to calculate p-values. p>0.05, ns; p≤0.05, *; p≤0.01,**; p≤0.001, ***. Data shown as mean±SD with individual data points.

To navigate this, we conducted a fluorescence assay with N-Methylmesoporphyrin IX (NMM), a small molecule probe specific for parallel G-quadruplexes^14,15^ (Fig. 1A and 1B), which indicated linM1 having a higher G4 forming propensity than circM1 at 100 mM salt concentration (Fig. 1E).

### Reverse Transcriptase Stalling confirms the presence of G-quadruplex

To further validate the presence of the G-quadruplex structure within the full-length M1 sequence, we performed a reverse transcriptase (RT) stalling assay. In the presence of K^+^, the sequence is expected to fold into a G4 structure, thereby impeding RT progression during cDNA synthesis. In contrast, under Li^+^ conditions—where G4 formation is unfavorable—the enzyme should extend cDNA without interruption^16^ (Fig. 2A). A primer complementary to the region immediately downstream of the predicted G-quadruplex was used for reverse transcription with SuperScript III, and the resulting cDNA products were amplified by PCR to quantify RT stalling associated with G4 formation.

**Figure 2.**
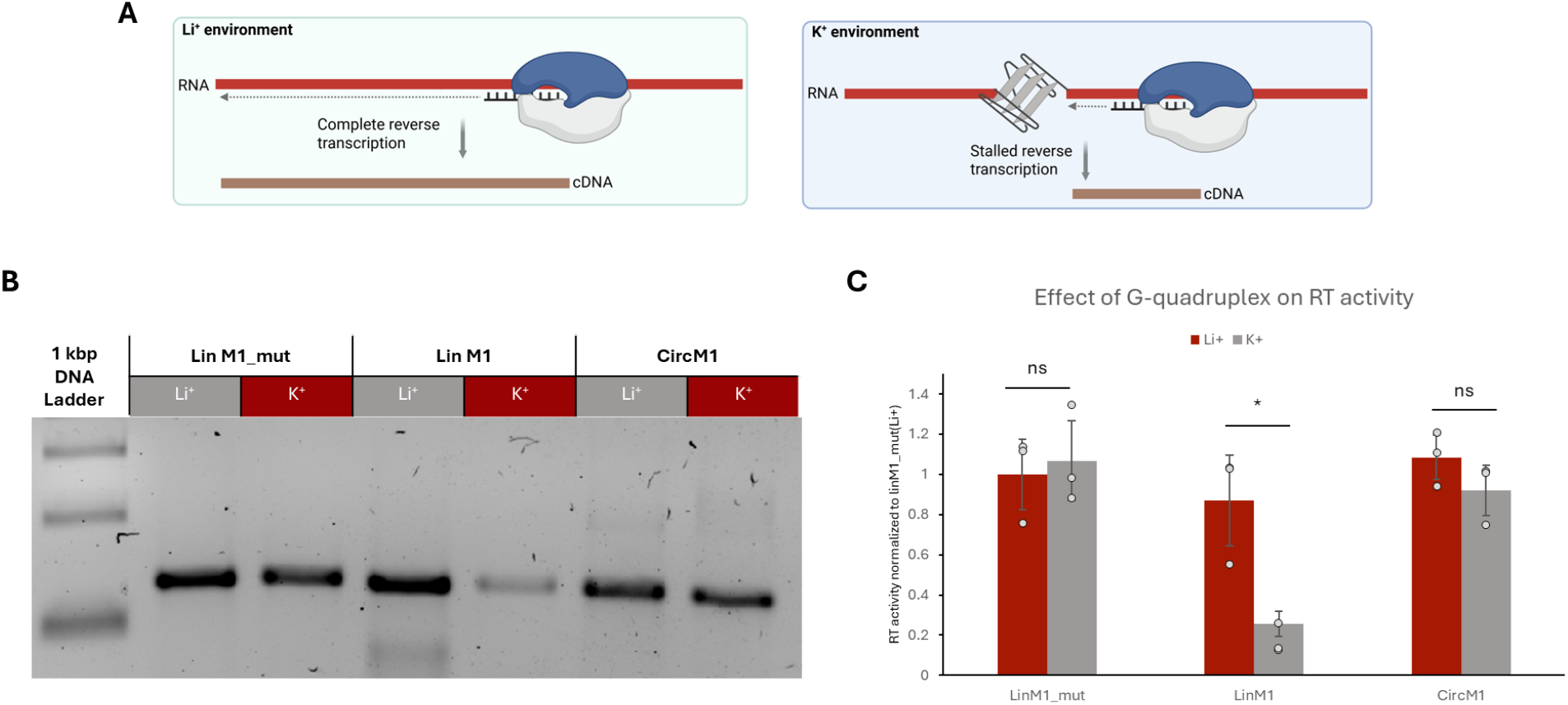
G-quadruplex effectively stalls reverse transcriptase in linM1. (A) Schematic representing the stalling of cDNA synthesis in K^+^ environment and unstalled reverse transcription in non-G4 promoting Li^+^ environment. (B) Agarose gel analysis showing amplification of the G4-containing region in mutated linM1, linM1 and circM1. (C) Quantitative analysis confirms the stalling of RT in linM1 by presence of G4. For quantitative analysis, n=3. Student’s t-test were performed to calculate p-values. p>0.05, ns; p≤0.05, *; p≤0.01,**; p≤0.001, ***. Data shown as mean±SD with individual data points.

A pronounced 70.5% reduction in reverse transcription activity was observed in K^+^ relative to Li^+^, confirming G-quadruplex formation within the M1 sequence. PCR amplification of cDNA generated under Li^+^ conditions also revealed a shorter band, which we attribute to RT skipping across the G4 region. To verify that this effect was G4-mediated, we mutated the region of interest using G4killer^17^ to disrupt G4 folding. In this mutant, no stalling was observed in K^+^ compared to Li^+^, further supporting that stalling in the wild-type sequence is G4 dependent.

Because our NMM assays indicated that circRNA has a reduced propensity to fold into G4 structures, we extended this assay to the circular form. Consistent with that observation, we did not detect a significant difference in RT activity between Li^+^ and K^+^ conditions for circRNA. Overall, these results demonstrate that the G4 of interest folds robustly in the full-length linear M1 RNA but has a much lower folding propensity in the circular form (Fig. 2B and 2C).

### Effect of G-quadruplex on M1 activity

Once we had established the higher propensity of linM1 to fold into a G-quadruplex, we then proceeded to study the effect of this tertiary structure on the catalytic activity of M1. Both linM1 and circM1 were allowed to fold in the presence of either Li^+^ or K^+^ as described earlier. The folded RNA was then added directly to the reaction buffer containing a well studied model pre-tRNA substrate, pATSerUG, labeled with Cy5 at the 5’ end and the reaction was conducted for 40 minutes (Fig. 3A). As expected, we observed a 7nt long leader as a product of the reaction. Interestingly, we saw a 21.5% decrease in activity of linM1 when allowed to fold in G-quadruplex favoring conditions. The activity of circM1 folded in K^+^ was only decreased to 10.1% compared with when folded in Li^+^ (Fig. 3B and 3C). This result is consistent with those of our previous assays indicating M1 isoforms are capable of forming G-quadruplexes and that linM1 has a higher propensity of G-quadruplex formation. More importantly, this suggests RNase P RNA can employ structural barriers to control their enzymatic activity. Furthermore, circM1 is resistant to such folding and is capable of performing its activity with little loss even in G-quadruplex favoring conditions. CircM1 might hence be crucial in maintaining pre-tRNA maturation in bacterial cells even in the presence of G-quadruplex forming environment for continued cellular processes and growth.

**Figure 3.**
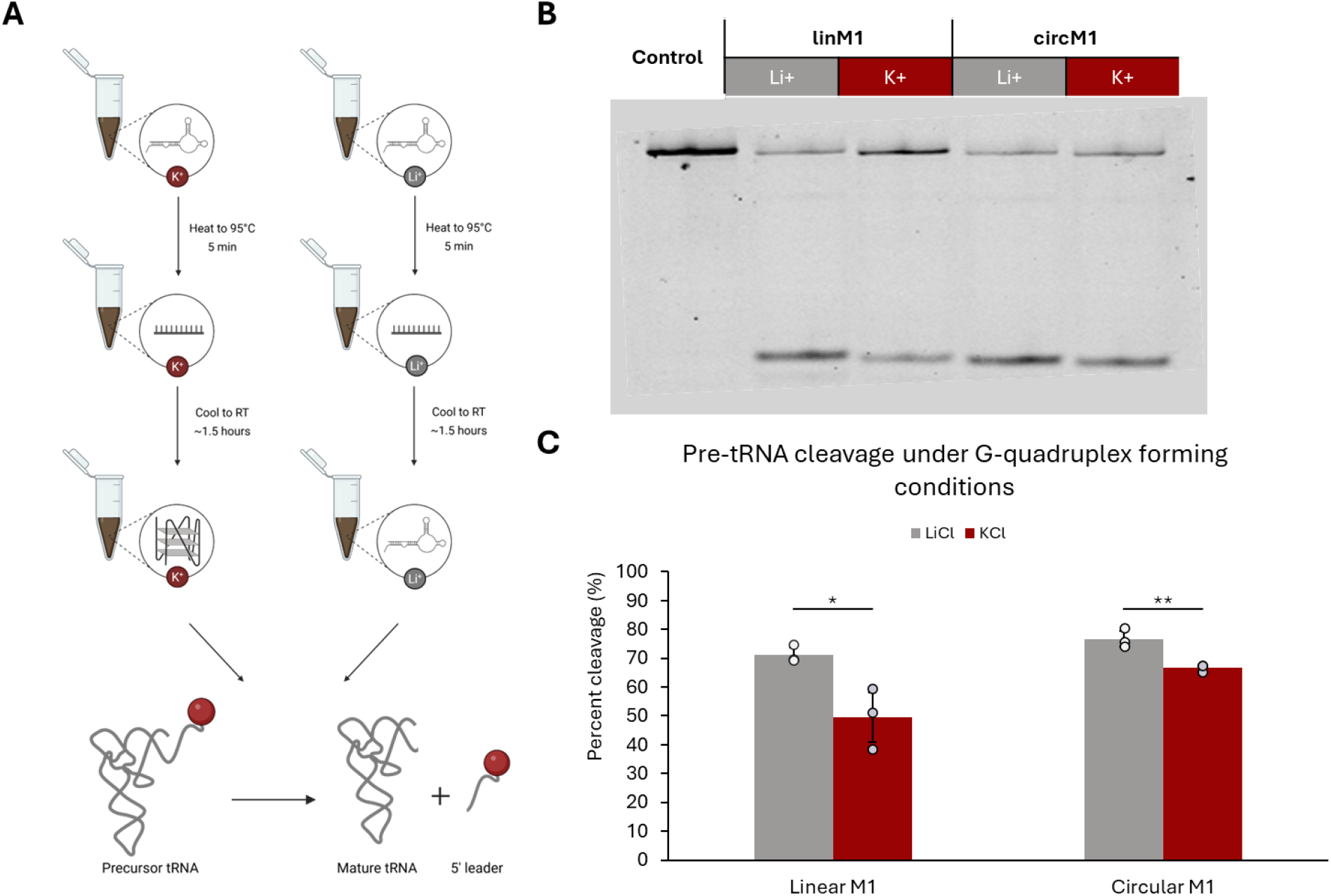
Catalytic activity of linM1 decreases in G4-stabilizing and non-stabilizing environments. (A) Schematic of pre-tRNA cleavage reactions to evaluate the effect of G4-formation on ribozyme efficiency. (B) Polyacrylamide gel displaying pre-tRNA substrate and products after cleavage reaction catalyzed by linM1 and circM1 in Li^+^ and K^+^ (C) Quantitative analysis confirms the partial loss of activity in G4 promoting conditions. LinM1 exhibits a higher loss compared to circM1. For quantitative analysis, n=3. Student’s t-test were performed to calculate p-values. p>0.05, ns; p≤0.05, *; p≤0.01,**; p≤0.001, ***. Data shown as mean±SD with individual data points.

### G4 potentially regulate M1 activity through a condensate mediated mechanism

To further understand the mechanism through which M1 activity might be affected due to this tertiary structure, we investigated the condensate properties of linM1 in G4 versus non-G4 favoring conditions by using Li^+^ and K^+^, respectively. It has been previously shown that M1, like most enzymes, catalyzes reactions in condensates^18^. Furthermore, various studies have investigated the role of G-quadruplexes in condensate formation^19,20^. By varying the salt concentration from 0 to 500 mM and condensate assisting protein, poly (L)-lysine (PLL) (Fig. 4A), to M1 RNA ratio from 0.1 to 5, we observed 5 distinct condensate architectures of Cy5 labeled M1 RNA (Fig. 4B). For equal charge ratios of RNA and protein, three salt conditions yielded distinct condensate behavior in Li^+^ versus K^+^, directly evidencing the role of G-quadruplex in condensation and thereby affecting the catalytic activity of M1 RNA isoforms (Fig. 4D). While it has been previously shown that G-quadruplexes promote condensate formation, we observed that under these three conditions, a less concentrated condensate architecture was favored in K^+^, leading to lower local concentrations of molecules and hence reduced activity of the M1 RNA.

**Figure 4.**
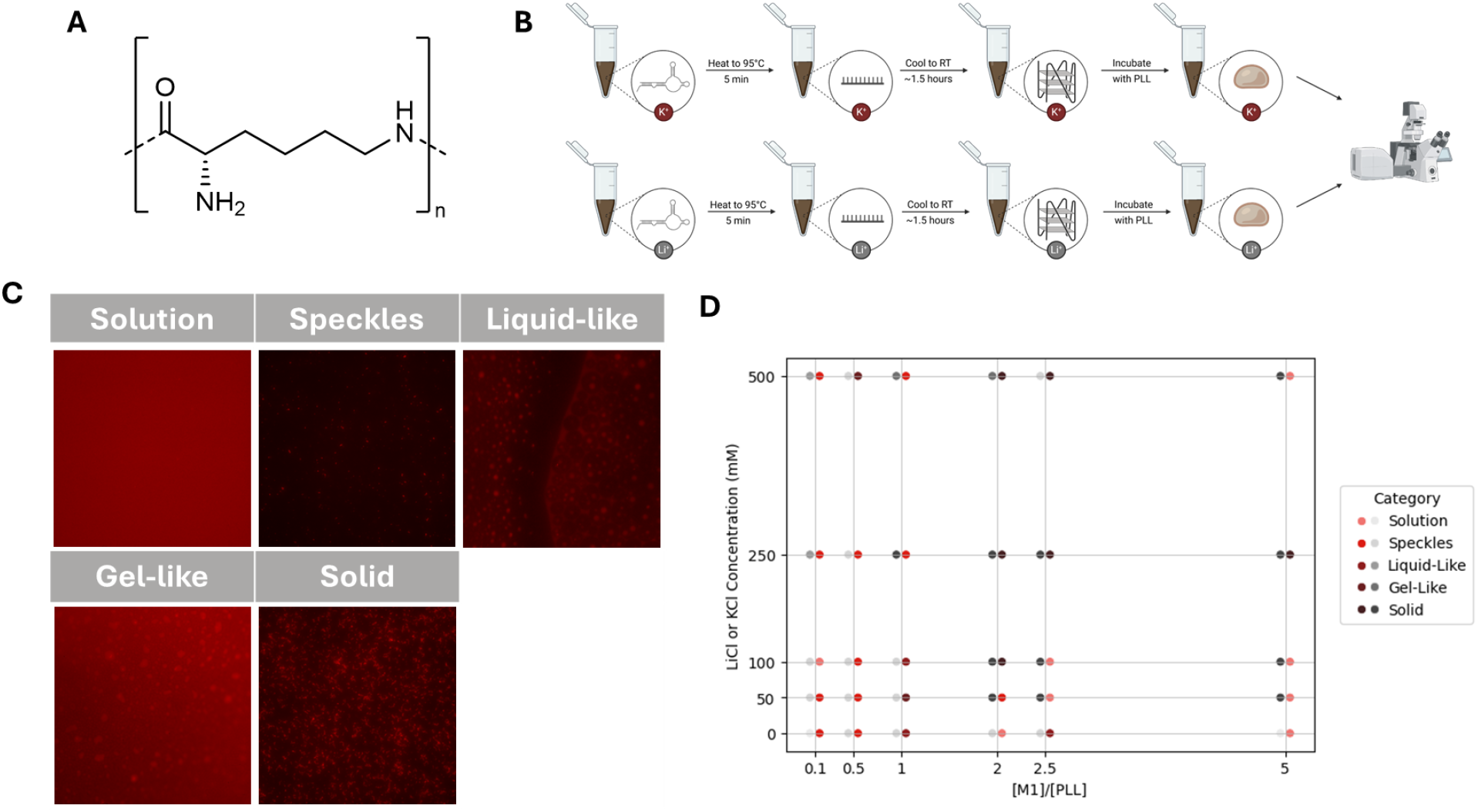
G4 formation affects condensate structure. (A) Chemical structure of poly-(L)-lysine (B) Schematic of M1-PLL condensate experiments (C) Fluorescence Microscopy images of the five distinct condensate architectures resulting from various M1:PLL ratios (0.1-5) and salt concentrations (0-500 mM) (D) Results of condensate experiments suggest G4 formation in linM1 lead to different condensate physical properties.

### RNase P RNA G4s are abundant across domains

RNase P is a universal enzyme and so, we investigated if G-quadruplex acts as a regulatory structure in RNase P RNA in other species as well. To address this, we obtained known and predicted sequences of RNase P RNA for 14,560 species through the Rfam database^21,22^. Pqsfinder predicted 28% of the sequences to contain putative G4 forming motifs (Fig. 5A). While we see a similar enrichment in bacterial class A, archaeal and eukaryotic RNase P, bacterial class B RNase P exhibits only 2.3% species containing putative G4 forming motifs (Fig. 5B). This can be attributed to their low GC-content and the protein playing a more enhanced role in enzyme stabilization, eliminating the need for a regulation through G4^23^. We then obtained the kingdom, phylum, class and orders of all the species through NCBI API^24^. We saw a normal distribution of RNase P RNA G4 containing species across all levels of classification and did not observe an enrichment (Fig. 5C). To further investigate the role of G4 in this RNA, we analyzed the bacterial class A species through BacDive^25^ and TEMPURA^26^. Again, no enrichment was seen for either gram stain (Fig. S5A), a pH dependency (Fig. S5B), growth temperature, maximum or minimum growth temperature (Fig. S5C and S5D). A limitation of this was that both BacDive and TEMPURA, while vast, are not complete and did not provide a complete picture of all the species. Additionally, no such databases exist which can be used for such specific analyses of archael and eukaryotic species, requiring further investigation into why G4 are formed in some species RNase P RNA but not others.

**Figure 5.**
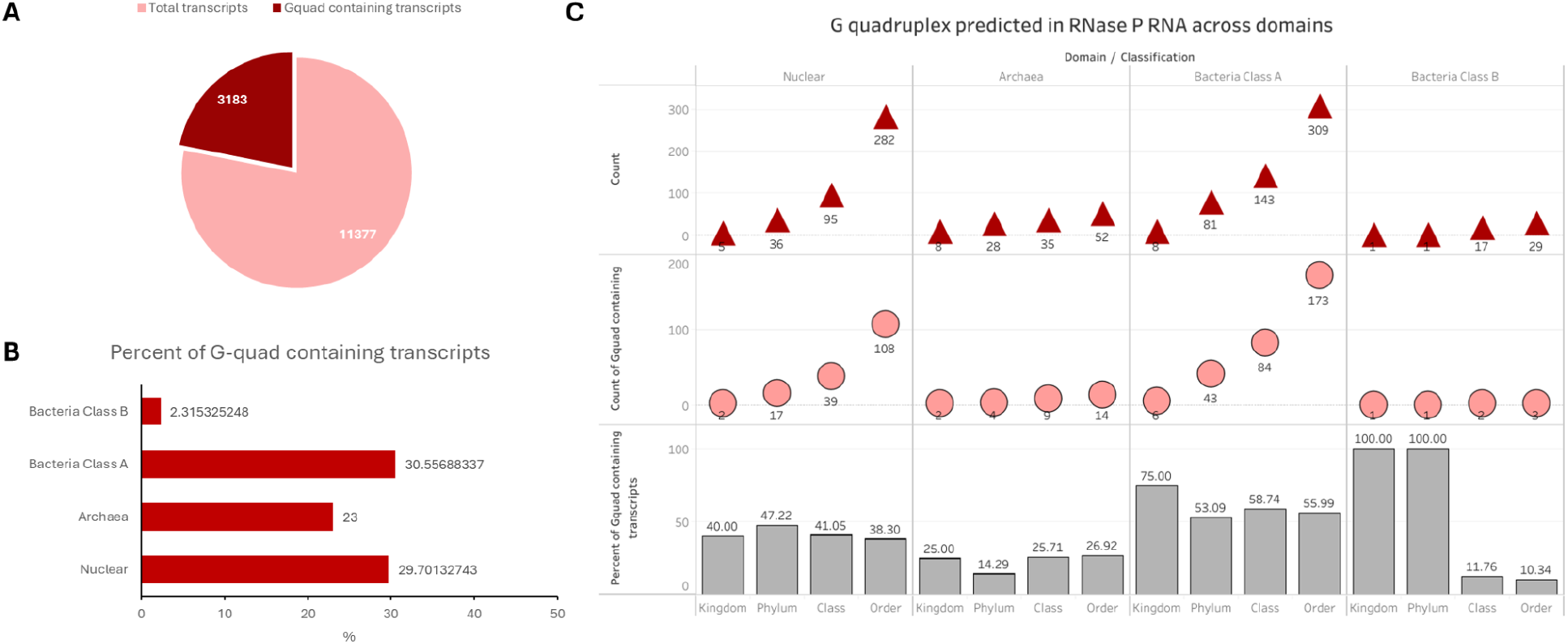
Global distribution and classification of predicted G-quadruplexes in RNase P RNA across domains of life. (A) Total transcripts analyzed versus the subset containing predicted G-quadruplexes (G4s), shown as a pie chart with counts of G4-positive and G4-negative transcripts. (B) Bar plots below summarize the percentage of G4-containing RNase P transcripts within each major domain/classification group, including Nuclear, Archaea, Bacterial Class A, and Bacterial Class B RNase P RNAs. (C) Domain-resolved visualization of predicted G-quadruplex abundance in RNase P RNA. Red triangles indicate total counts of predicted G4 motifs across taxonomic levels (kingdom, phylum, class, and order), while pink circles represent the number of unique G4-containing transcripts. Gray bars show the percentage of RNase P RNAs containing at least one predicted G4 motif within each taxonomic category. Together, these data illustrate that G-quadruplexes are present across all surveyed domains but vary widely in frequency and distribution, with notably lower prevalence in bacterial Class B RNase P RNAs compared to other groups.

## Conclusions

In this study, we identify the first functional RNA G-quadruplex in bacteria, embedded within the RNase P M1 RNA. We show that this G4 forms far more readily in the linear isoform than in its circular counterpart, and that its presence markedly suppresses linM1 catalytic activity, likely through a condensate-mediated mechanism. These findings reveal an unrecognized layer of structural regulation in bacterial RNase P and suggest that RNA G-quadruplexes may play broader roles in shaping bacterial RNA biology. Moving forward, determining whether additional predicted G4s within M1 RNA adopt similar structures and influence activity will be essential. Together, our results establish a foundation for exploring RNA G-quadruplexes as functional regulatory elements in bacteria.

## Supplementary Information

**Table S1.**
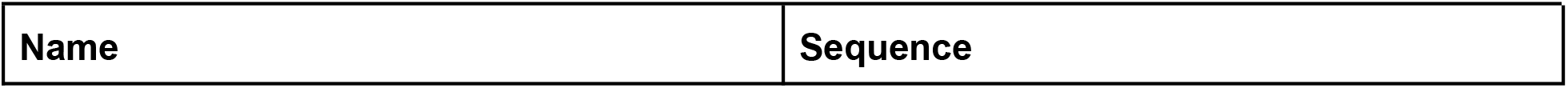
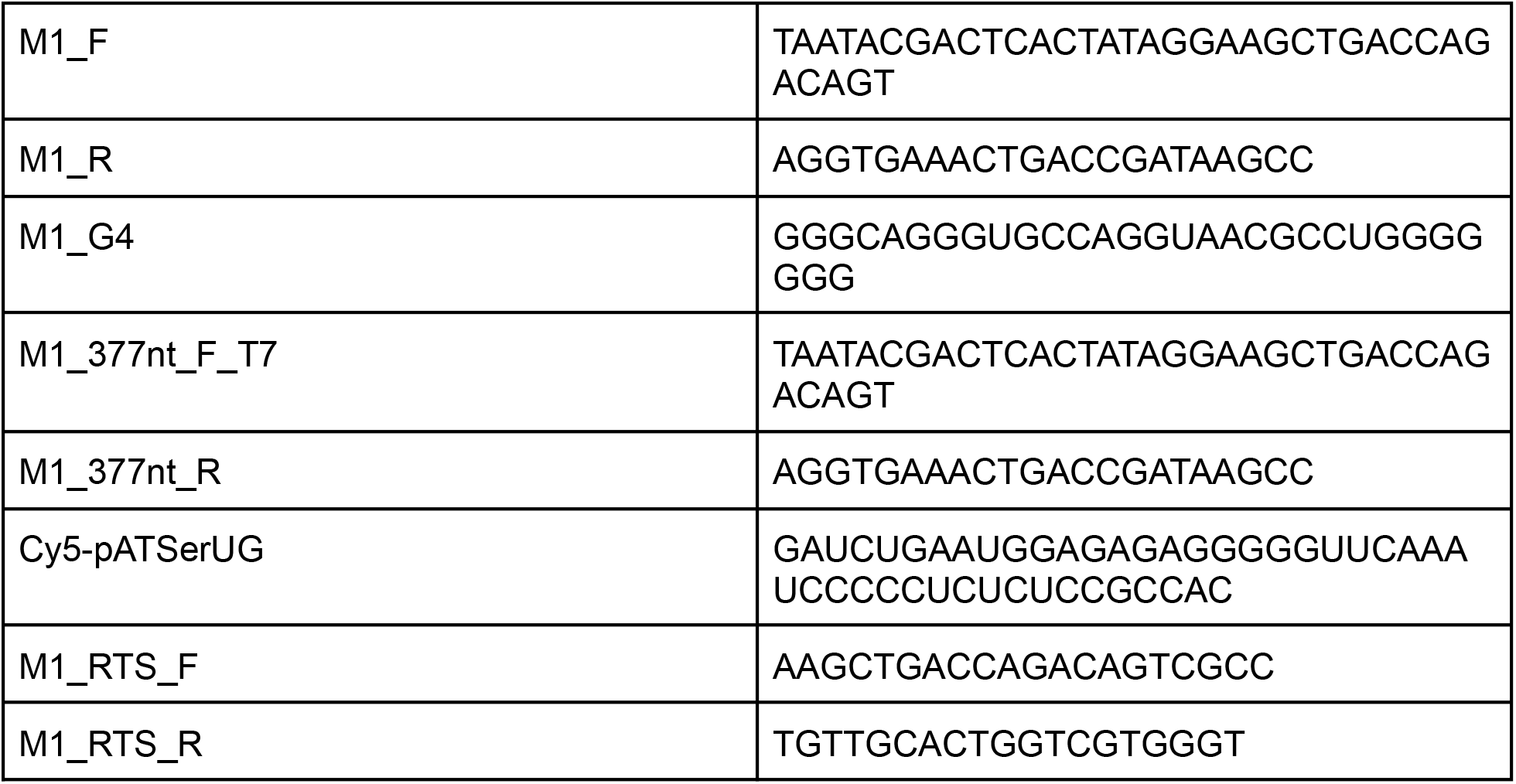
List of DNA and RNA oligonucleotides.

**Table S2.** List of chemical reagents.

| Name | Manufacturer |
| --- | --- |
| Poly-(L)-lysine Hydrobromide | Fisher Scientific |
| N-methyl mesoporphyrin IX (NMM) | Fisher Scientific |
| Cy5-Adenosine triphosphate | TriLink |
| Cacodylic acid, 98% | Fisher Scientific |
| Potassium chloride | Invitrogen |
| Lithium chloride | Sigma Aldrich |
| Superscript III | Invitrogen |
| MgCl<sub>2</sub> | Invitrogen |
| Trizma hydrochloride solution pH 7.5 | Sigma-Aldrich |
| dNTP mix (10 mM) | Invitrogen |
| Dithiothreitol (0.1 mM) | Invitrogen |

**Table S3.** List of instruments.

| Name | Manufacturer |
| --- | --- |
| Mermade 4 | LGC Biosearch Technologies |
| LSM 880 Confocal Microscope | Zeiss |
| Gel Imager | BioRad |
| Quant Studio 5 qPCR | Applied Biosystems |

## Methods

### Synthesis of Cy5-pre-tRNA

Model pre-tRNA, pATSerUG, was synthesized using solid-support oligonucleotide synthesis at 1 μM scale with DMT-ON using Mermade 4 synthesizer. The RNA was cleaved from the bead support and bases were deprotected in a single step using 1100 μL 1:1 ammonia/methylamine by shaking at room temperature for 4 hours. This solution was filtered using Pierce columns and the beads were washed twice with 300 μL water. Ammonia was evaporated under vacuum and the solution was lyophilized to obtain 2’ protected DMT-ON RNA. This was dissolved in 125 μL of DMSO and incubated in 60 μL of TEA and 75 μL of TEA/H3F to deprotect the 2’ hydroxyls. RNA was then purified and DMT was cleaved off using RNA purification cartridge. The obtained RNA solution was lyophilized and redissolved in nuclease free water. The obtained RNA was analyzed using 12% denaturing polyacrylamide gel.

### Circular Dichroism

In a 1 mL reaction volume, 1 μM RNA in 20 mM lithium cacodylate buffer and 100 mM of LiCl or KCl was heated at 95°C for 5 minutes and allowed to cool to room temperature at the rate of 1°C/minute. The solution was then transferred to a quartz cuvette. CD spectra were recorded in the range of 220 nm to 340 nm at 25°C. Five readings were recorded and averaged using Jasco J-1500 CD spectrophotometer.

### NMM fluorescence assay

In a 20 μL reaction volume, 1 μM RNA in 20 mM lithium cacodylate buffer and 100 mM of LiCl or KCl was heated at 95°C for 5 minutes and allowed to cool to 25°C at the rate of 1°C/minute. NMM was added to a final concentration of 2.5 μM and the mixture was incubated in the dark for 10 minutes. Fluorescence intensities were recorded using an excitation wavelength and emission wavelength of 399 nm and 605 nm, respectively.

### Pre-tRNA cleavage assay

In a 2 μL reaction volume, 1 μM RNA in 20 mM lithium cacodylate buffer and 100 mM of LiCl or KCl was heated at 95°C for 5 minutes and allowed to cool to room temperature at the rate of 1°C/minute.

0.48uL of the above prepared solution was to the reaction buffer (40mM MgCl2, 50mM Tris HCl pH 7.2, 5% PEG8000, 100mM NH4Cl) followed by addition of 0.05μM pATSerUG. The reaction (2μL) was incubated at 37°C for 40 minutes and quenched by addition of 8μL of 2X OrangeG in formamide. The solution was denatured at 90°C for 2 minutes and quickly cooled on ice for 1 minute followed by loading on a 20% denaturing polyacrylamide gel. Quantitation of gel bands was performed using Bio-rad ImageLab software. p-values were calculated using student’s t-test.

### Condensate formation with PLL

In a 2 μL reaction volume, 0.1-5 μM RNA in 20 mM lithium cacodylate buffer and 100 mM of LiCl or KCl was heated at 95°C for 5 minutes and allowed to cool to room temperature at the rate of 1°C/minute. To this, 2 μL of an equal charge ratio of PLL in 20 mM lithium cacodylate buffer and 100 mM of LiCl or KCl was added, vortexed, centrifuged and transferred to a black-bottom 96-well plate and analyzed using Leica microscope using a 40X air objective. Images were processed with FIJI.

### M1, Cy5-M1 and circM1 synthesis

DNA templates corresponding to the required RNA sequences with T7 promoter sequence at the 5’ end were purchased from TWIST Biosciences, amplified using PCR and purified using DNA Clean & Concentrator-5. MEGAshortscript T7 Transcription Kit was used for RNA synthesis. 375 mM Guanosine Monophosphate (GMP) was added to the M1 synthetic reaction. This allowed the incorporation of monophosphate at the 5’ end. 0.5mM Cy5 labeled CTP was added to the reaction for synthesis of Cy5-M1. The reaction was carried out at 37°C for 15 hours followed by DNase I treatment at 37°C for 15 minutes. The RNA was precipitated overnight, centrifuged, washed and air-dried as described above. M1 RNA was then analyzed using a 6% denaturing polyacrylamide gel.

For circularization, M1 RNA was circularized using T4 RNA Ligase I and purified using a 6% denaturing polyacrylamide gel. The gel pieces were incubated in 0.3M NaOAc and 0.02 mg/mL glycogen and shaken overnight at 4°C. The solution was filtered through a 0.2-micron filter to remove the gel pieces and 3X volume of 100% ethanol was added to the filtrate. This was incubated at -80°C overnight, centrifuged and washed as described earlier.

### Reverse transcriptase stalling assay

Method was adapted from Kwok et al (2016). Briefly, 3pmol of M1 RNA, 1uM of M1_RTS_R and nuclease free water were mixed to a total volume of 5.5uL and incubated at 75°C for 3 minutes and immediately placed on ice. To this, 3 uL of reaction buffer was added for final concentrations of 150 mM LiCl, 4 mM MgCl2, 20 mM Tris-HCl pH 7.5, 1 mM DTT, and 0.5 mM dNTPs. The reaction mixture was incubated at 35°C for 5 minutes, followed by addition of 0.5uL of SuperScript III and incubation of 15 minutes at 50°C. 5 uL of the cDNA was used for a 12.5uL PCR reaction as described in the manufacturer’s protocol. Products were analyzed on a 2% agarose gel and quantitatively analyzed using BioRad Image Lab.

**Figure S1.**
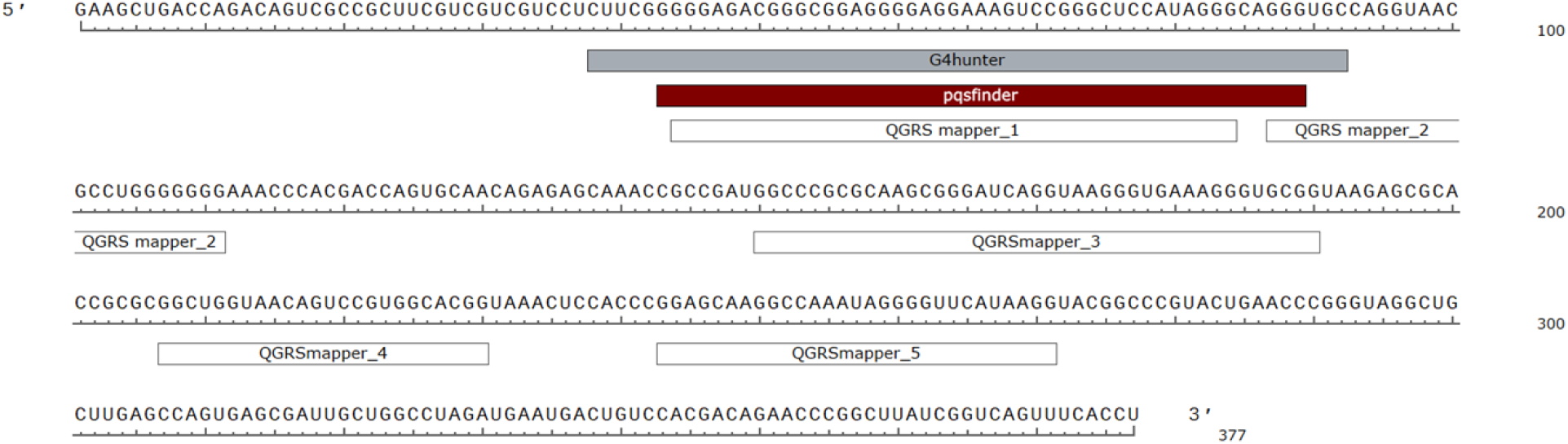
G-quadruplexes predicted using three different tools—pqsfinder, G4hunter and QGRS mapper

**Figure S2.**
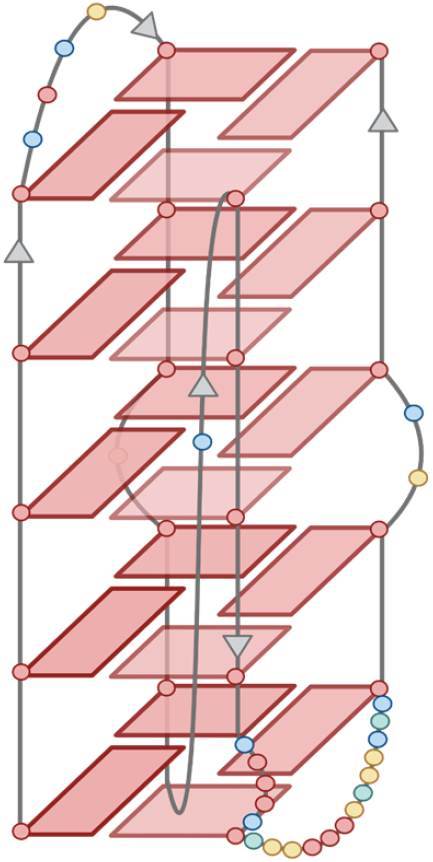
Structure of the 47nt long predicted G-quadruplex in M1 RNA consisting of 5 tetrated and 3 bulges.

**Figure S3.**
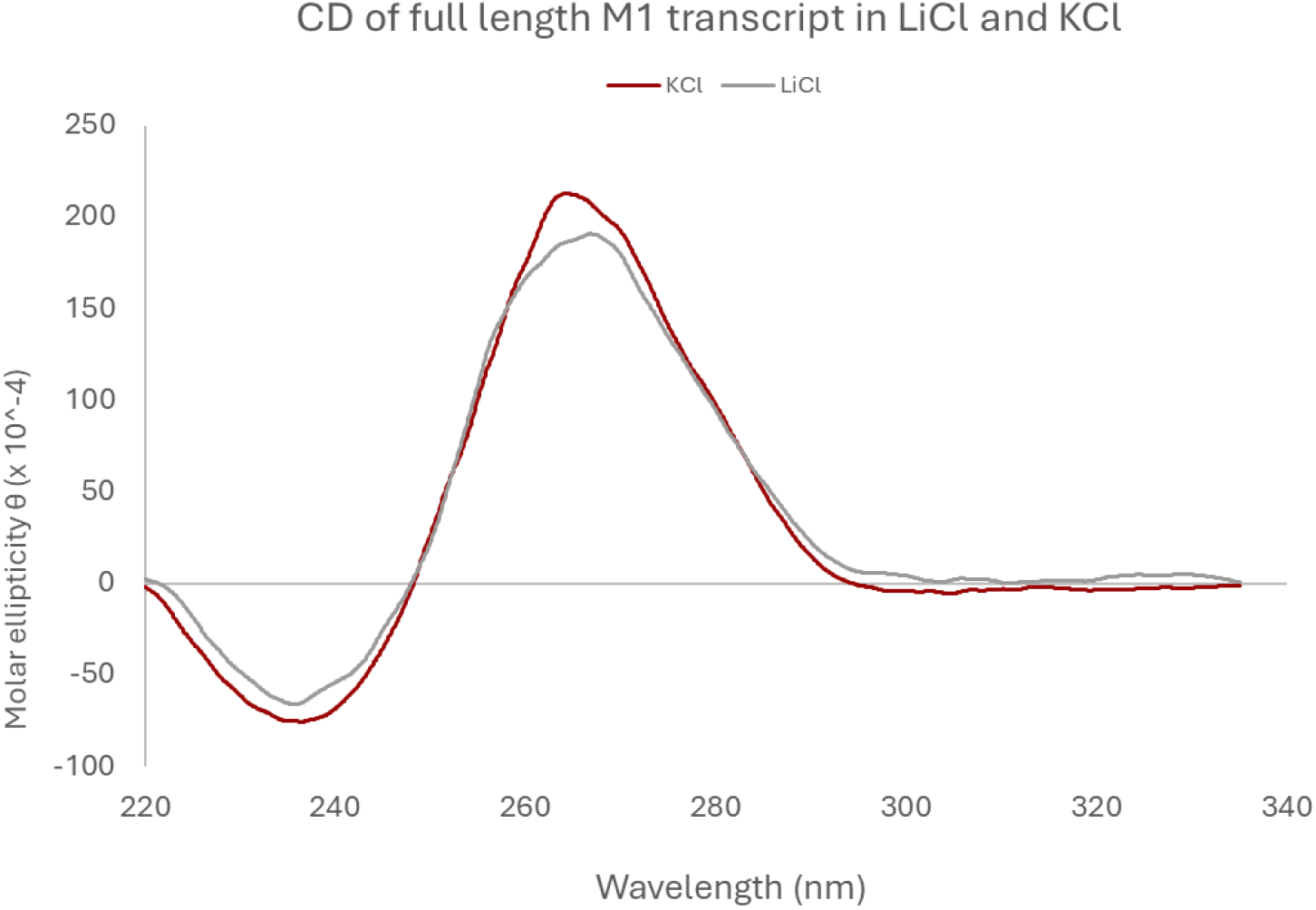
CD spectra of 377nt long M1 RNA in 100mM LiCl or KCl

**Figure S4.**
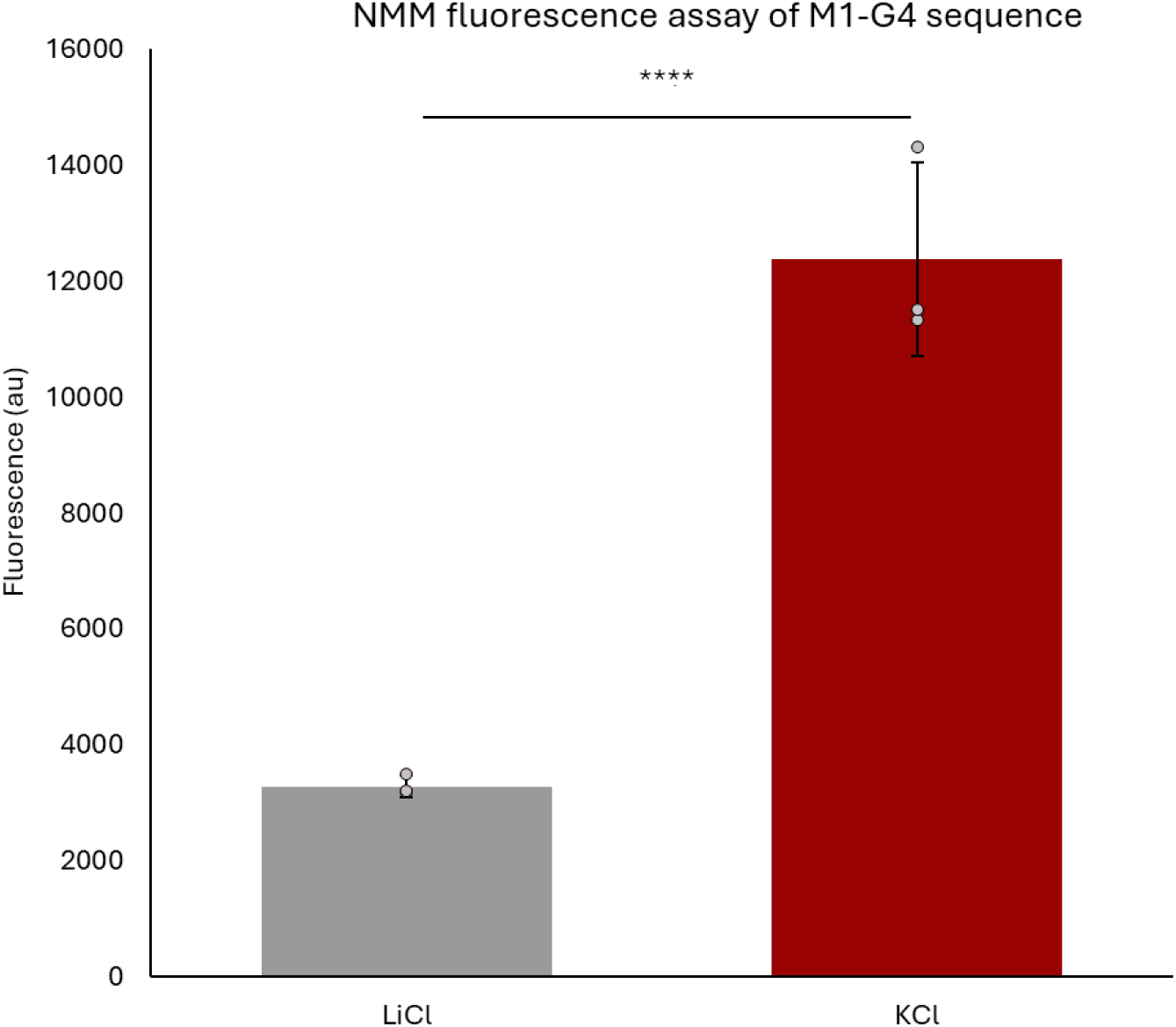
NMM fluorescence of 47nt putative G4 M1 RNA fragment in 100mM LiCl or KCl. For quantitative analysis, n=3. Student’s t-test were performed to calculate p-values. p>0.05, ns; p≤0.05, *; p≤0.01,**; p≤0.001, ***.

**Figure S5.**
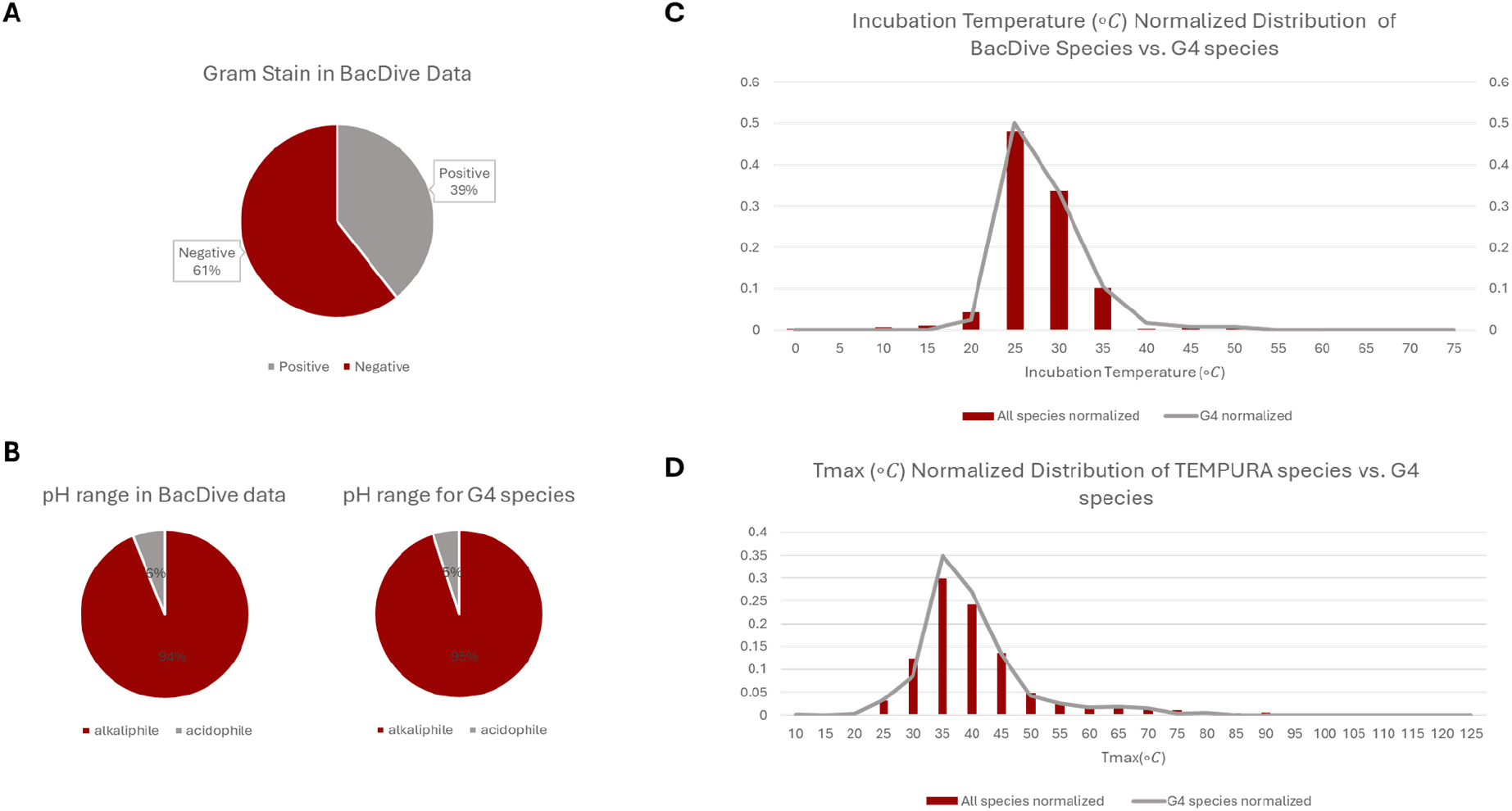
G4 forming bacterial class B RNase P RNA were not found to be enriched for a specific physiological trait. (A) Gram stain enrichment analysis of G4 forming RNase P RNA (B) pH range enrichment analysis of G4 forming RNase P RNA (C) Incubation temperature enrichment analysis of G4 forming RNase P RNA (D) Maximum growth temperature enrichment analysis of G4 forming RNase P RNA.

